# StarBEAST2 brings faster species tree inference and accurate estimates of substitution rates

**DOI:** 10.1101/070169

**Authors:** Huw A. Ogilvie, Remco R. Bouckaert, Alexei J. Drummond

## Abstract

Fully Bayesian multispecies coalescent (MSC) methods like *BEAST estimate species trees from multiple sequence alignments. Today thousands of genes can be sequenced for a given study, but using that many genes with *BEAST is intractably slow. An alternative is to use heuristic methods which compromise accuracy or completeness in return for speed. A common heuristic is concatenation, which assumes that the evolutionary history of each gene tree is identical to the species tree. This is an inconsistent estimator of species tree topology, a worse estimator of divergence times, and induces spurious substitution rate variation when incomplete lineage sorting is present. Another class of heuristics directly motivated by the MSC avoids many of the pitfalls of concatenation but cannot be used to estimate divergence times. To enable fuller use of available data and more accurate inference of species tree topologies, divergence times, and substitution rates, we have developed a new version of *BEAST called StarBEAST2. To improve convergence rates we add analytical integration of population sizes, novel MCMC operators and other optimisations. Computational performance improved by 13.5× to 13.8× when analysing empirical data sets, and an average of 33.1 × across 30 simulated data sets. To enable accurate estimates of per-species substitution rates we introduce species tree relaxed clocks, and show that StarBEAST2 is a more powerful and robust estimator of rate variation than concatenation. StarBEAST2 is available through the BEAUTi package manager in BEAST 2.4 and above.

## 1 Introduction

The throughput of sequencing technologies has improved many-fold over the past two decades culminating in next generation sequencing (NGS), and it is now feasible to sequence whole or partial genomes or transcriptomes for phylogenetic studies (Lemmon and Lemmon, 2013). NGS produces hundreds or thousands of phylogenetically useful loci (see for example Blom *et al*., 2016) with potentially millions of sites spread across a data set of multiple sequence alignments.

While NGS offers hundreds or thousands of loci at relatively low cost, making accurate inferences from the enormous amount of data produced is particularly challenging. In the case of *BEAST, a fully Bayesian method of species tree inference which implements a realistic and robust evolutionary model in the multispecies coalescent (MSC; Degnan and Rosenberg, 2009; Heled and Drummond, 2010), it becomes exponentially slower as the number of loci in an analysis is increased. This scaling behaviour causes *BEAST to become intractably slow after a certain number of loci (the exact number will depend on other parameters of the data set, see Ogilvie *et al*., 2016). Given the current challenges of using large phylogenomic data sets with *BEAST there have been three broad alternatives available to researchers; concatenate sequences from multiple loci, use heuristic methods statistically consistent with the MSC, or choose a tractable subset of loci to use with a fully Bayesian method like *BEAST, BEST (Liu, 2008), or BPP (Yang, 2015).

Using maximum likelihood phylogenetic methods to infer a species tree based on concatenated sequences will return the single tree that best fits the combined sequence alignment according to the phylogenetic likelihood function (Felsenstein, 1981). Popular maximum-likelihood concatenation methods include RAxML, PAML and PhyML (Stamatakis, 2014; Yang, 2007; Guindon *et al*., 2010). Bayesian methods, such as ExaBayes and BEAST (Aberer *et al*., 2014; Drummond and Rambaut, 2007), will instead return a distribution of trees which are probable given the combined sequence alignment, a set of priors, and the same likelihood function. Recent results show that likelihood-based concatenation can be counterproductive, producing statistically inconsistent results which assign high confidence to incorrect nodes due to model misspecification (Liu *et al*., 2015). In the so-called “anomaly zone” of short branch lengths, the most probable gene tree topology will be different from the species tree, and estimated tree topologies will likely differ from the true species tree topologies (Degnan and Rosenberg, 2006; Kubatko and Degnan, 2007).

More recently identified problems with likelihood-based concatenation are systematic errors when estimating branch lengths, including overestimation of divergence times. Because some time is required for genes to coalesce looking backwards from a speciation event, the expected molecular distance between two species is greater than if coalescent events occurred simultaneously when gene flow ceases. This leads concatenation to overestimate the divergence times across a species tree in proportion to effective population size (Arbogast *et al*., 2002; Ogilvie *et al*., 2016).

Incomplete lineage sorting (ILS) also causes systematic errors in estimated branch lengths when using concatenation. When a gene tree topology is discordant with the species tree topology, then the gene tree will contain one or more branches that define splits not occurring in the species tree. If a substitution occurs on one of these discordant gene tree branches, the resulting site pattern would define a homoplasy on the species tree, implying multiple substitutions. This effect has been termed substitutions produced by ILS (SPILS) and causes concatenation to overestimate the lengths of specific branches and underestimate the lengths of others, which produces apparent substitution rate variation where none exists (Mendes and Hahn, 2016). For all the above reasons, trees inferred using concatenation are therefore not a reliable approximation of the species tree in terms of branch lengths or topology.

As an alternative to concatenation for use with phylogenomic data, heuristic methods which do not perform phylogenetic likelihood calculations but are statistically consistent with the MSC have been developed. These include summary methods which utilise distributions of estimated gene tree topologies as input, such as the rooted triplet method MP-EST (Liu *et al*., 2010) and the quartet method ASTRAL (Mirarab *et al*., 2014a). Another quartet method is SVDquartets, which utilises single-nucleotide polymorphism (SNP) matrices (Chifman and Kubatko, 2014). Recent results show that MP-EST should be used with caution as it is sensitive to gene tree errors (Mirarab and Warnow, 2015; Xi *et al*., 2015). At low levels of ILS, MP-EST is less accurate than likelihood-based or neighbour-joining concatenation at inferring topologies, and even at high levels of ILS it may be no more accurate than concatenation (Ogilvie *et al*., 2016). No available heuristic method is both statistically consistent and can infer branch lengths in substitution or calendar units. Therefore heuristic methods cannot be reliably used to estimate divergence times. If concatenation is used to estimate branch lengths or divergence times for a species tree topology estimated by another heuristic method, then those estimates will be unreliable for the same reasons as pure concatenation.

An issue specific to summary methods is when the assumption of no recombination within loci is frequently violated because overly long loci are used (Gatesy and Springer, 2013). To resolve larger and deeper species trees using summary methods, longer and more informative loci may be required to infer more accurate gene trees. However the larger and deeper a tree, the more recombination events will have occurred. The use of longer loci and the higher incidence of recombination will both increase the risk of recombination occurring within loci, which has been dubbed the “recombination ratchet” (Springer and Gatesy, 2016).

As an alternative to increasing locus length, fully Bayesian MSC methods like *BEAST infer more accurate gene trees by sharing information between loci through the species tree (Szöllősi *et al*., 2015). In this way very accurate species trees can be estimated using only weakly informative loci, which may not be possible using MP-EST (Xu and Yang, 2016). To avoid the recombination ratchet summary methods can use naïvely binned subsets of gene trees estimated by *BEAST (Zimmermann *et al*., 2014), or statistically binned subsets of genes trees estimated by concatenation (Mirarab *et al*., 2014b). Statistical binning has been criticised as statistically inconsistent (Liu and Edwards, 2015), and for either binning method the resulting species trees still cannot be used for molecular dating.

With the aim of improving the computational performance of fully Bayesian MSC inference of species trees, we have developed an upgrade to *BEAST — StarBEAST2 — which is available as a package for BEAST 2 (Bouckaert *et al*., 2014). By improving computational performance StarBEAST2 enables the use of more loci, which will improve the precision of estimated parameters and provide an alternative to concatenation. We have also developed and include in StarBEAST2 new MSC relaxed clock models to enable accurate inference of per-species substitution rates.

## 2 New Approaches

### 2.1 Analytical integration of population sizes

Markov Chain Monte Carlo (MCMC) methods like *BEAST jointly integrate over many parameters by proposing small changes at each step to eventually produce a probability distribution for all parameters. From a researcher’s perspective, some may be “nuisance” parameters not of scientific interest. For example species tree topology and divergence times may be of interest, but not effective population sizes. For tractable parameters, an analytic solution will integrate over the entire range of values at each MCMC step, and may be faster than MCMC integration. However explicit estimates will not be produced so this approach is suitable only for nuisance parameters. Among-site rate variation is already integrated out at each step; the likelihood of each site is calculated for all possible discrete gamma rates at each step, so individual site rates are not estimated (Yang, 1994).

Analytical integration of constant per-branch population sizes was first implemented as part of BEST (Liu *et al*., 2008), and is described in detail by Jones (2017). The analytic solution, which we have added to StarBEAST2, uses an inverse gamma conjugate prior for population sizes. By default StarBEAST2 fixes the shape of the distribution *α* = 3 and only estimates the mean of the distribution *μ*, which is proportional to the scale parameter *β*:

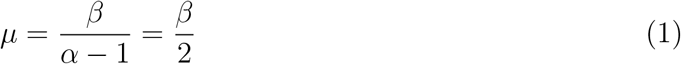

In this special case where *α* = 3, the standard deviation is identical to the mean:

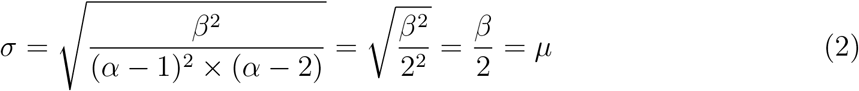

The coefficient of variation *C*_v_ = σ/*μ* of the prior distribution for effective population sizes is therefore 1.

### 2.2 Coordinated tree topology changing operators

One approach to improving the performance of MSC analyses which simultaneously estimate gene and species trees (such as *BEAST) is to develop MCMC operators which propose coordinated changes to both the species tree and the gene trees in the same step. Yang and Rannala (2014) introduced a Metropolis-Hastings (MH; Metropolis *et al*., 1953; Hastings, 1970) operator which makes nearest-neighbour interchange (NNI) changes to the species tree topology, and simultaneously makes changes to gene tree topologies which preserve compatibility of the gene trees within the proposed species tree. Later, both Jones (2017) and Rannala and Yang (2017) introduced more general coordinated operators which make subtree prune and regraft (SPR) changes to the species tree. We have reimplemented these coordinated NNI and SPR moves in StarBEAST2 as a single new operator called “CoordinatedExchange”. Rannala and Yang (2017) also describe a proposal distribution which favours topological changes on shorter branches as well as less radical changes in topology. StarBEAST2 implements a simpler proposal distribution but still favours less radical changes by applying adjustable proposal probability weights to (less radical) NNI moves and (more radical) SPR moves.

### 2.3 Coordinated node height changing operators

A novel class of coordinated Metropolis operators was introduced by Jones (2017), which pick at random a non-root non-leaf species tree node *S* with an existing height of *t*(*S*). A new height *t*′(*S*) is chosen from a uniform distribution with lower and upper bounds D and U. The height of the species tree node and the heights of subtrees of gene tree nodes (termed “connected components”) are all shifted by the amount η = *t*′(*S*) − *t*(*S*).

The *D* and *U* bounds limit the minimum and maximum values of *η* to those which do not require modifying the topology of the gene tree or of the species tree, and the algorithm to determine those bounds is given by Jones (2017). The species tree root node is excluded because there is no natural upper bound in that case. As long as the connected components are chosen with reference only to the topology of the species tree, the topology of the gene trees, and the mapping of sampled individuals to species, operators of this class are symmetric.

We have developed a new operator called “CoordinatedUniform” that belongs to this general class but has not been implemented before. Individuals from extant species which descend from a species tree node, or are directly descended from a gene tree node, are referred to here as “descendant individuals”. The gene tree nodes *s* selected by this operator to be shifted in height are all those for which:

1. at least one descendant individual of *s* is also a descendant individual of the *left* child of *S*
2. at least one descendant individual of *s* is also a descendant individual of the right child of *S*
3. all descendent individuals of *s* are also descendent individuals of *S*

An example of how gene tree nodes are selected and node heights shifted is given in Supplementary Material.

We have also developed a new adaptive MH (Andrieu and Thoms, 2008) operator called “CoordinatedExponential” which changes the height of the species tree root node and the height of connected components by an amount *η*. The gene trees nodes to be shifted are chosen using identical criteria as for CoordinatedUniform. Because this operator changes the height of the root node, a different method must be used to pick *η* compared to CoordinatedUniform.

First the lower bound *D* is identified in the same way as for CoordinatedUniform and as described in Jones (2017). The difference between *D* and the current root height is referred to as *x*, and a new random value *x*′ is chosen from an exponential distribution. The value of *x*′ − *x* is then used for *η*. The median of the exponential distribution is adaptively modified over the course of an MCMC chain to equal the posterior expectation of *x*.

Because the proposal distribution for a new species tree root height is independent of the current height, the Hastings ratio which is usually *q*(*x*′,*x*)/*q*(*x*, *x*′) (Hastings, 1970) can be simplified to *π*(*x*)/*π*(*x*′). The natural logarithm of the Hastings ratio may then be derived from the respective probability densities of *x* and *x*′ drawn from an exponential distribution with the rate λ:

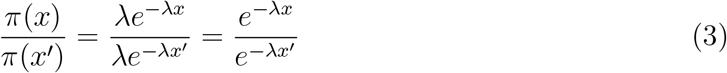

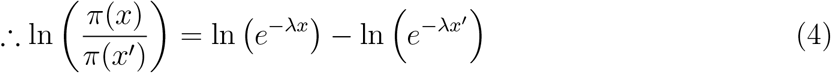

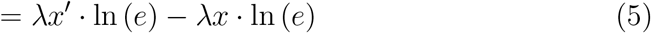

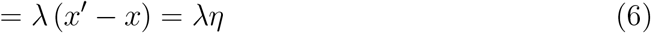

### 2.4 Species tree relaxed clocks

The overall rate of evolution occurring at a given locus within a species will be influenced by the nature of the particular gene and also by the natural history of the particular species. For a given gene, the average substitution rate may depend on the effects of selection such as the accelerated molecular evolution of sex-biased genes in *Arabidopsis thaliana* (Gossmann *et al*., 2014), and on within-genome variation in mutation rate (Baer *et al*., 2007). For a given species, the average substitution rate is correlated with a multitude of traits including metabolic rate, body size, and fecundity, although causal relationships are difficult to pin down (Bromham, 2011). Unsurprisingly in light of the above, empirical analysis has shown that two major factors contributing to rate variation among gene branches are the per-gene rate and the per-species rate (Rasmussen and Kellis, 2007).

Because variation is expected in the nature of different genes and species, and therefore variation is also expected in the average substitution rate of different genes and species, multispecies coalescent models should take both per-gene and per-species rate variation into account. *BEAST can accommodate both types of rate variation using gene tree relaxed clock models (for examples see Berv and Prum, 2014; Lambert *et al*., 2015). This involves estimating per-branch substitution rates separately for each branch of each gene tree. While gene tree relaxed clocks may accommodate variation in substitution rates between species, they do not produce estimates of species branch rates. To enable accurate inference of species branch rates, we have developed a new species tree relaxed clock model.

The challenge of applying a relaxed clock to the species tree is that phylogenetic likelihood calculations require branch rates for each branch of each gene tree. Our clock model computes those rates using the total expected number of substitutions 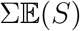 accumulated along the entire length of a gene tree branch. Substitutions are expected to be accumulated at the mean clock rate of the gene tree *c*, multiplied by the lengths of time *L* spent traversing each species tree branch, multiplied by the rates *R* of the corresponding species tree branches. Typical nuclear substitution rates for mammals are around 10^−3^ substitutions per site per million years (Phillips *et al*., 2009).

The gene tree branch rates r can then be derived by dividing the total expected number of substitutions by the total length of that branch *l*. The gene tree branch rates for the illustrated example (Figure 1; Table 1) are therefore:

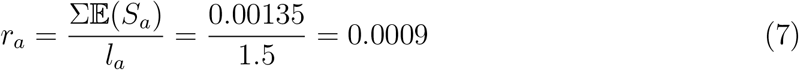

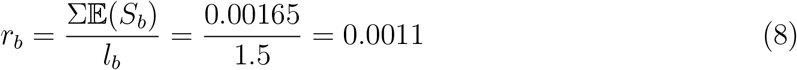

**Figure 1:**
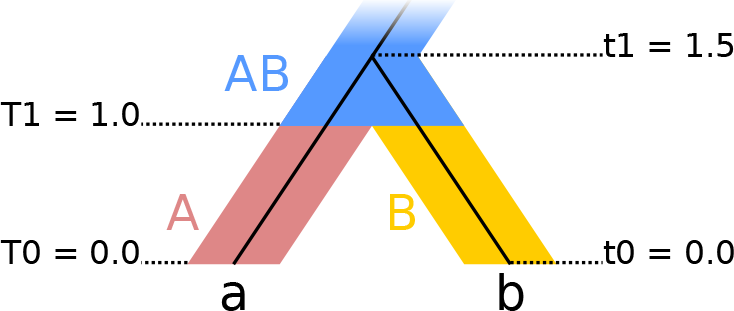
Two-species phylogeny used to illustrate species tree relaxed clocks. There are two extant species “A” and “B”, and one ancestral species “AB”. Within the species tree there is a single gene tree with extant individuals “a” and “b”. The single speciation event occurs at time T1, and the single coalescence event occurs at time t1. Gene tree rates are computed according to Table 1.

**Table 1:** Expected number of substitutions 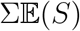 for gene branches *a, b* under a species tree relaxed clock

| Gene branch | Gene rate<sup>1</sup> $c$ | Length<sup>2</sup> $L$ | | | Species rate<sup>3</sup> $R$ | | | $\mathbb{E}(S) = c \cdot L \cdot R$ | | | $\Sigma\mathbb{E}(S)$ |
| --- | --- | --- | --- | --- | --- | --- | --- | --- | --- | --- | --- |
|  |  | A | B | AB | A | B | AB | A | B | AB |  |
| a | 0.001 | 1.0 | 0.0 | 0.5 | 0.7 | 1.0 | 1.3 | 0.00070 | 0.00000 | 0.00065 | 0.00135 |
| b |  | 0.0 | 1.0 | 0.5 |  |  |  | 0.00000 | 0.00100 | 0.00065 | 0.00165 |

<sup>1</sup> The overall substitution rate at a given locus.
<sup>2</sup> The length of a given gene tree branch within species tree branch A, B or AB.
<sup>3</sup> The substitution rate of species tree branch A, B or AB.

The new species tree relaxed clock model is available in StarBEAST2. Branch rate models that can be used with a species tree relaxed clock currently include the well-established uncorrelated log-normal (UCLN) and uncorrelated exponential (UCED) models (Drummond *et al*., 2006), as well as the newer random local clock model (Drummond and Suchard, 2010).

The current species tree relaxed clock implementation estimates—separately for each species tree branch—a single relative rate. However it is possible to imagine a further relaxed model that estimates—again separately for each species tree branch—hyperparameters for a prior distribution on substitution rates. This would enable gene tree branch rates to be guided by the species tree, but still allow some difference in response between genes.

## 3 Results and Discussion

### 3.1 StarBEAST2 correctly implements the multispecies coalescent

New methods must be shown to be correct implementations of the target model. One way to accomplish this for MCMC methods is to estimate parameters from a prior distribution using the MCMC kernel, and to also draw independent samples from the same distribution by simulation. The resulting parameter distributions should be identical if the implementation is correct. We used this method to test the correctness of the novel features in StarBEAST2; analytical population size integration, coordinated operators, and species tree relaxed clocks. Simulated and StarBEAST2 distributions were identical for species and gene tree topologies (Figure S1,S2), species and gene tree node heights (Figure S3,S4), and for gene tree branch rates (Figure S5,S6). This combination of results supports the correctness of the StarBEAST2 implementation.

### 3.2 Species tree relaxed clocks prevent SPILS

When using concatenation to infer a species tree, SPILS causes apparent substitution rate variation. However in an ultrametric (time tree) framework like BEAST, branch lengths are constrained so that terminal species begin at time zero. We hypothesised that if a relaxed clock is used with concatenation in an ultrametric framework, SPILS will be absorbed as faster substitution rates for lineages that would be lengthened by SPILS in a non-ultrametric framework.

In an ultrametric framework with a strict clock and no external (e.g. fossil, biogeographical or known clock rate) calibrations, the substitution rate of each branch is set to 1. This ensures that 1 unit of time is equivalent to 1 expected substitution. Using a relaxed clock with no external calibrations the substitution rate of each branch can vary, but the expectation of the mean rate of all branches is 1, preserving the relationship of 1 unit of time = 1 expected substitution. Therefore when SPILS causes the rates of some branches to be faster than 1, the rates of some other branches will be slower than 1 to keep the expected mean constant.

We used BEAST concatenation and StarBEAST2 with a species tree relaxed clock to infer the branch lengths and substitution rates of simulated species trees with the topology ((((A,B),C),D),E), using sequence alignments simulated using a strict clock. Gene tree discordance will increase the estimated length of A, B and C branches for these species trees (Mendes and Hahn, 2016), and as hypothesised substitution rates for A and B branches inferred using concatenation were biased towards being faster than the true rate of 1 (Figure 2). Estimated substitution rates for the C branch were more variable, and could be faster or slower than 1. Substitution rates estimated for the D and E branches were biased towards being slower than 1, presumably to balance the mean rate. Concatenation also overestimated the lengths of tip branches, another known bias when using concatenation to infer a species tree (Ogilvie *et al*., 2016). No biases were observed for the branch rates or lengths estimated using StarBEAST2 (Figure 2).

**Figure 2:**
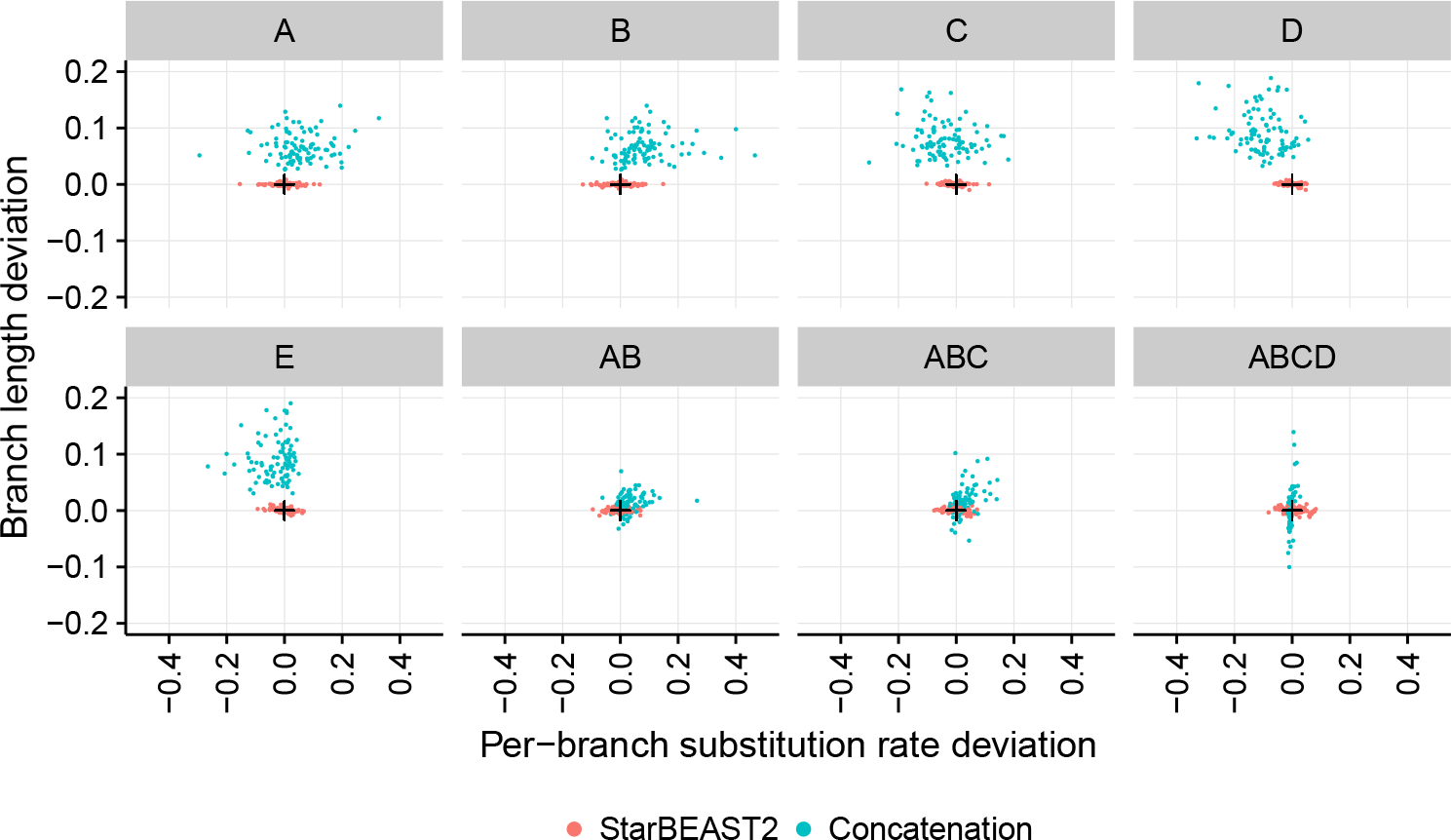
Accuracy of branch substitution rates and lengths inferred by BEAST concatenation and StarBEAST2. Deviation is the difference of each estimated rate and length from the true value. Estimated rates and lengths are the posterior expectation of the overall substitution rate and length for each species tree branch. Black crosses in each panel indicate the point of perfect accuracy. Each panel shows the distributions for the labelled extant or ancestral branch. N = 96.

A number of estimated branch rates had 95% credible intervals that excluded the true rate of 1 when using concatenation. If a study is testing whether substitution rates vary across a species tree, those branch rates could be erroneously interpreted as faster or slower than average. In our simulations, the clock rate of the D branch would be inferred as slower than average in 37 out of 96 replicates (Figure S7), despite the sequence data being simulated using a strict clock. When applying the same 95% credible intervals to branch lengths, the true simulated length was excluded with just two exceptions for all tip branches across all replicates using concatenation (Figure S8). In contrast, no erroneous results would be inferred for branch rates given the same data using StarBEAST2, and out of the 768 total simulated non-root branch lengths, only five erroneous results would be inferred (Figure S7,S8).

Mendes and Hahn (2016) demonstrated that SPILS causes systematic bias when estimating branch lengths, and we show that this translates into systematic bias when estimating per-branch substitution rates. Because the bias is caused by ILS which is a function of population sizes and branch lengths, there is no reason to expect that large trees with varying population sizes and branch lengths would be any less biased.

### 3.3 Datasets used to characterise the new methodological approaches

To characterise the performance of coordinated operators, methods of population size integration and relaxed clocks, we tested StarBEAST2 using empirical and simulated sequence data. The empirical data set used for this analysis is from the North American chorus frog genus Pseudacris, and was originally collected and analysed by Barrow *et al*. (2014). This data set has sequences from 26 nuclear loci across 44 sampled individuals. The individuals belong to 19 extant Pseudacris lineages and two outgroup species. Barrow *et al*. (2014) reported phased haplotypes but to avoid wasting computational resources we used a single haplotype per individual.

A key metric of phylogenies that can be used to judge whether it is necessary to employ MSC models is the average branch length in coalescent units *τ*(2*N*_*e*_)^−1^. In this study Ne will always refer to the effective population size of diploid individuals. Given short branch lengths, likelihood-based or neighbour-joining concatenation is unable to infer accurate species trees regardless of the number of loci used, but for long branch lengths, concatenation is approximately as accurate as *BEAST (Ogilvie *et al*., 2016). Indeed concatenation can be considered a special case of the MSC as the models converge when gene trees are identical to the species tree (Liu *et al*., 2015). Using StarBEAST2, the average branch length within this genus was estimated to be 2.81*τ*(2*N*_*e*_)^−1^. This is an intermediate average length compared to the shallow simulations analysed by Ogilvie *et al*. (2016) which had a shorter average length of 1.08*τ*(2*N_e_*)^−1^.

Each replicate of each *Pseudacris* empirical analysis used the same sequence data, and the true species tree topology, dates and rates were not known with certainty. For performance results more generally applicable than a single empirical system, and to measure the coverage and accuracy of StarBEAST2 inference, we created a simulated data set of 30 replicates. A unique species tree was simulated for each replicate, and gene trees and locus sequences were simulated according to the MSC.

We simulated 26 nuclear loci from 21 extant species with two individual haplotypes per species, very similar to the empirical data set size. The simulation parameters, including the birth rate, death rate and population sizes, were also chosen to be similar to estimated *Pseudacris* parameters. The simulated data set had an average branch length of 2.99*τ*(2*N*_e_)^−1^, so the relative accuracy of MSC models compared to concatenation should be comparable with empirical systems like *Pseudacris*.

### 3.4 Coordinated height changing operators and analytical integration improve performance

To determine which configuration of new features would achieve the best performance, we ran StarBEAST2 using different combinations of operators, methods of population size integration and clock models. To measure convergence both effective sample size (ESS) per hour and ESS per million states were computed for each independent chain. ESS per hour can be used to calculate the total time required for a converged chain (nominally where ESS equals or exceeds 200), and reflects how effectively operators explore the space of trees and parameters, as well as the computational time required by each operator proposal and likelihood calculation. In contrast, ESS per million states reflects only the exploration of tree and parameter space independently of calculation times. A variety of statistics were recorded for each analysis (Table S2-S7), and for each replicate the slowest ESS rate out of all statistics recorded for that individual chain was used for all subsequent analyses.

Multiple linear regressions with log transformed ESS rates as the response variables were used to measure the effect of coordinated topology changing operators, coordinated node height changing operators, and the method of population size integration. Each additional feature was treated as a binary indicator variable so that we could quantify the relative performance as a percentage by exponentiating the coefficient for each addition (Table 2).

**Table 2:**
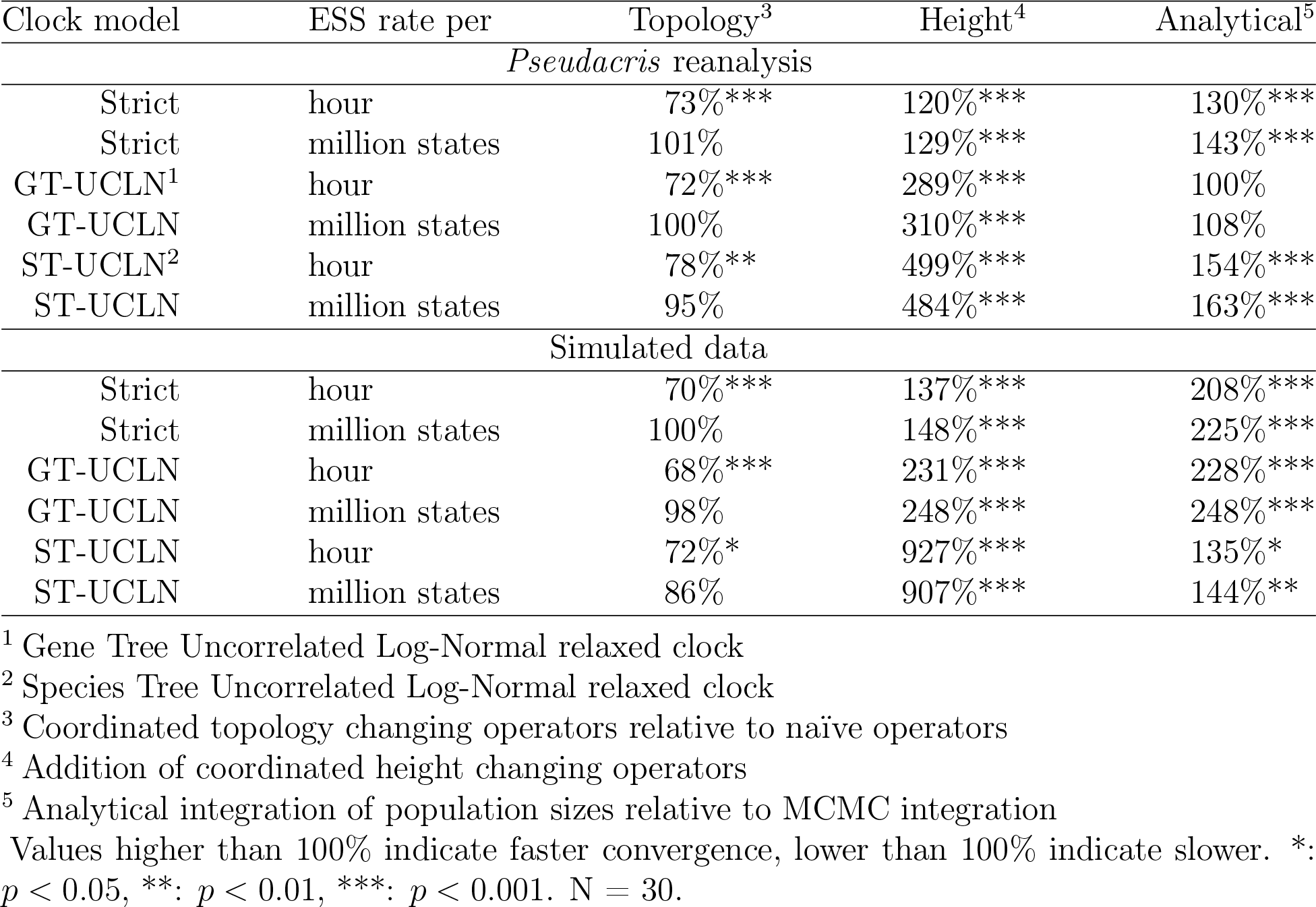
Relative performance of operators, population size integration and clock models.

| Clock model | ESS rate per | Topology<sup>3</sup> | Height<sup>4</sup> | Analytical<sup>5</sup> |
| --- | --- | --- | --- | --- |
| *Pseudacris* reanalysis |  |  |  |  |
| Strict | hour | 73%*** | 120%*** | 130%*** |
| Strict | million states | 101% | 129%*** | 143%*** |
| GT-UCLN<sup>1</sup> | hour | 72%*** | 289%*** | 100% |
| GT-UCLN | million states | 100% | 310%*** | 108% |
| ST-UCLN<sup>2</sup> | hour | 78%** | 499%*** | 154%*** |
| ST-UCLN | million states | 95% | 484%*** | 163%*** |
| Simulated data |  |  |  |  |
| Strict | hour | 70%*** | 137%*** | 208%*** |
| Strict | million states | 100% | 148%*** | 225%*** |
| GT-UCLN | hour | 68%*** | 231%*** | 228%*** |
| GT-UCLN | million states | 98% | 248%*** | 248%*** |
| ST-UCLN | hour | 72%* | 927%*** | 135%* |
| ST-UCLN | million states | 86% | 907%*** | 144%** |

<sup>1</sup> Gene Tree Uncorrelated Log-Normal relaxed clock
<sup>2</sup> Species Tree Uncorrelated Log-Normal relaxed clock
<sup>3</sup> Coordinated topology changing operators relative to naïve operators
<sup>4</sup> Addition of coordinated height changing operators
<sup>5</sup> Analytical integration of population sizes relative to MCMC integration
Values higher than 100% indicate faster convergence, lower than 100% indicate slower. \*: $p < 0.05$ , \*\*: $p < 0.01$ , \*\*\*: $p < 0.001$ . $N = 30$ .

Coordinated topology operators consistently and significantly reduced ESS per hour, but had no significant effect on ESS per million states (Table 2), suggesting that coordinated topology operators are no more effective than naïve operators at proposing new states. A decrease in the number of states per hour (Figure S9) shows that they are more computationally expensive than naïve operators, and explains the negative effect on ESS per hour.

Coordinated height changing operators consistently and significantly increased ESS per hour and per million states, however the degree of improvement depended on the clock model (Figure 3). For strict clock analyses the increase in ESS per hour was modest at 1.2 times and 1.37 times for empirical and simulated data respectively, whereas for species tree relaxed clocks the increase was 4.99 times and 9.27 times respectively (Table 2). The difference in species tree relaxed clock performance suggests that coordinated height changing operators are necessary for practical implementations of that model.

**Figure 3:**
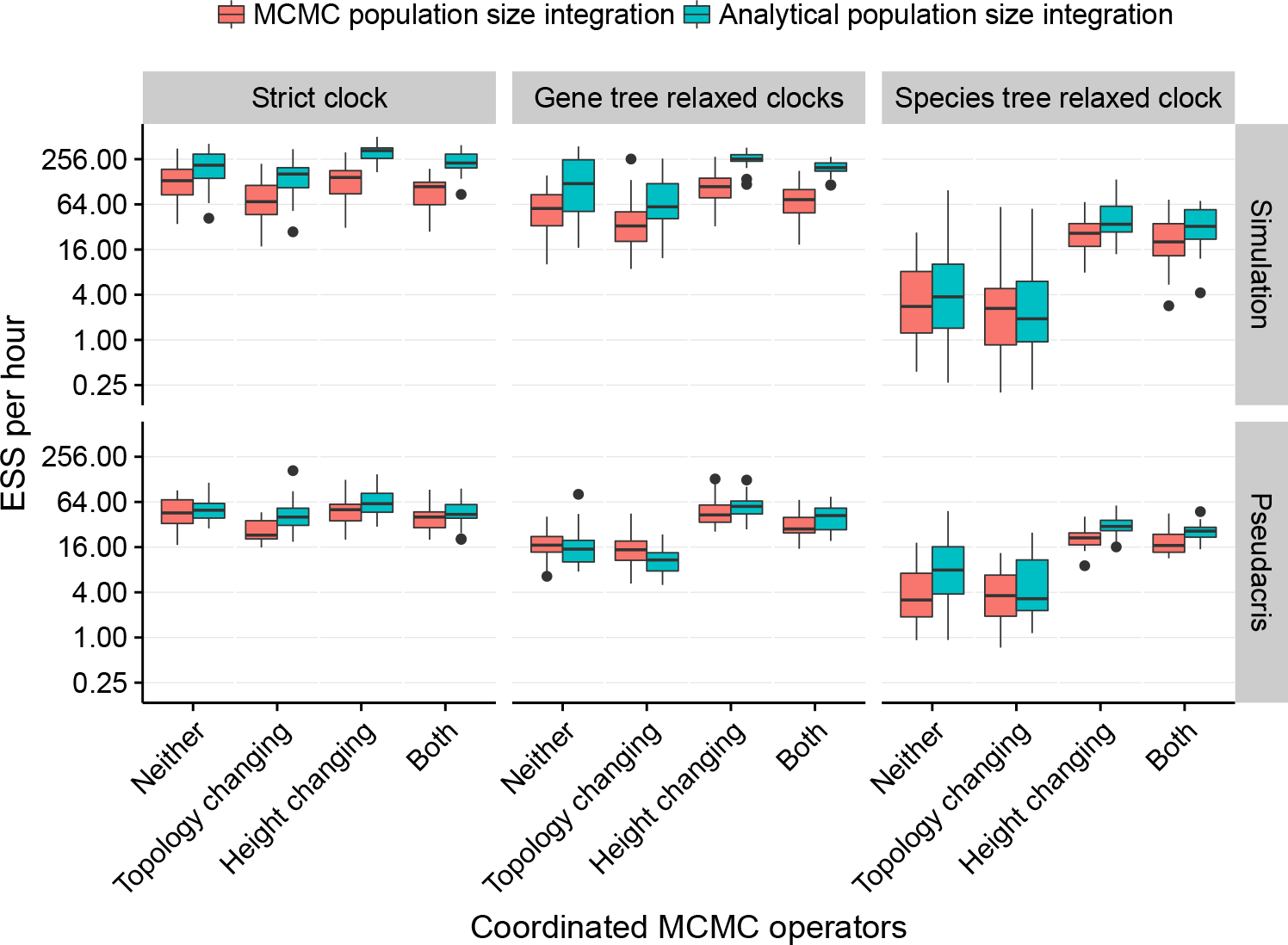
Impact of operators, population size integration and clock models on convergence. The estimated sample size (ESS) per hour for a given replicate used the smallest ESS out of all recorded statistics. Topology refers to the replacement of naïve nearest-neighbour interchange and subtree prune and regraft operators with coordinated operators. Height refers to the addition of operators which make coordinated changes to node heights. Uncorrelated log-normal relaxed clocks were applied to each gene tree (GT-UCLN) or to the species tree (ST-UCLN). N = 30.

Analytical population size integration significantly improved ESS per hour and per million states performance in all cases, with the exception of gene tree relaxed clocks applied to the *Pseudacris* data set (Table 2).

Even with new operators and analytical population size integration, the ESS per hour rates for species tree relaxed clocks were slower than for other clock models (Figure 3). One reason is that changing a species tree branch rate requires updating the phylogenetic likelihood for all gene trees, so the computational cost is much higher than for strict or gene tree relaxed clocks (Figure S9).

### 3.5 StarBEAST2 is an order of magnitude faster than *BEAST

StarBEAST2 also optimises the core multispecies coalescent algorithms by caching intermediate values and by using fast data structures. Operator weights have also been refined by manual iteration for better performance. Building on our results, by default StarBEAST2 enables coordinated height changing operators and analytical population size integration, but keeps naïve topology operators. To measure the combined improvement when StarBEAST2 is applied to *Pseudacris* data we compared the performance of StarBEAST2 with default settings to *BEAST. For the simulation data set, we compare StarBEAST2 with *BEAST and also with concatenation.

NGS data sets may have hundreds or thousands of loci. To gauge the performance of StarBEAST2 applied to these data sets we tested an empirical NGS data set; ultraconserved element (UCE; Faircloth *et al*., 2012) sequences from Philippine shrews of the genus Crocidura (Giarla and Esselstyn, 2015). This data set consists of 1112 loci sampled from a total of 19 individuals, which belong to 9 extant lineages. Again multiple statistics were recorded to compute ESS rates for each replicate.

Our simulation study confirmed that StarBEAST2 is many times faster than *BEAST (Figure 4). For simulated data the average log convergence rate of StarBEAST2 with gene tree relaxed clocks was 5.54 ln(^*ESS*^/*hour*). This compares to 2.04 using *BEAST, an increase in performance of exp(5.54 − 2.04) = 33.1 times (Table S2). In fact, StarBEAST2 was exp(4.18 − 2.04) = 8.5 times faster at analysing 52 loci than *BEAST was when analysing 26.

**Figure 4:**
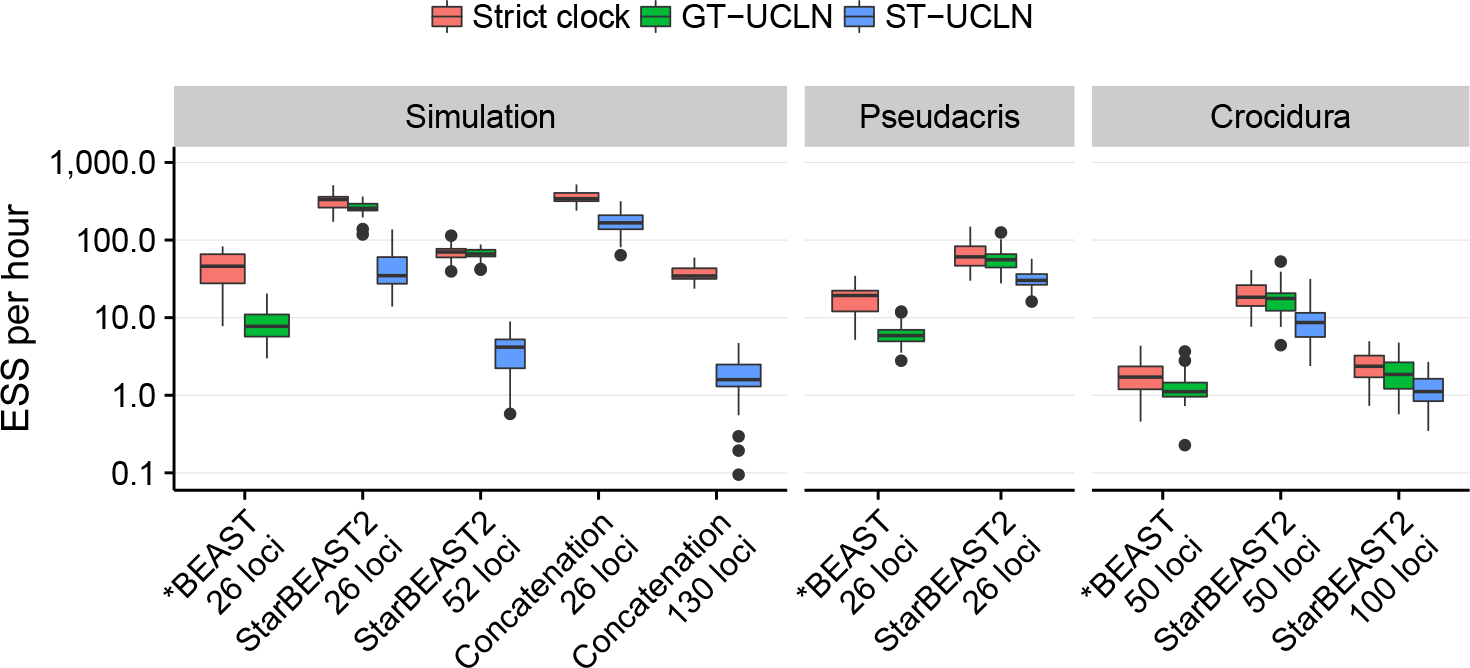
Convergence of different methods applied to simulated and empirical data sets. The estimated sample size (ESS) per hour for a given replicate used the slowest ESS rate out of all recorded statistics. Methods are BEAST concatenation, *BEAST, and StarBEAST2 with uncorrelated log-normal relaxed clocks applied to each gene tree (GT-UCLN) or to the species tree (ST-UCLN). Two *Pseudacris* *BEAST outliers with ESS rates below 0.1 are not shown. N = 30.

StarBEAST2 was an order of magnitude faster when analysing either empirical data set. For gene tree relaxed clock reanalyses of *Pseudacris* the difference was exp(3.98 −1.38) = 13.5 times. For 50-locus *Crocidura* reanalyses it was exp(2.79 − 0.17) = 13.8 times (Table S4,S6).

The ESS per hour convergence of species tree relaxed clocks was lower than for gene tree relaxed clocks. When applying StarBEAST2 to simulated data, gene tree relaxed clocks were exp(5.54 − 3.71) = 6.2 times faster than using species tree relaxed clocks (Table S2). The difference was much smaller for empirical data; for *Pseudacris* reanalyses gene tree relaxed clocks were exp(3.98−3.44) = 1.7 times faster, and for *Crocidura* they were exp(2.79−2.12) = 2.0 times faster (Table S4,S6). In all three cases species tree relaxed clocks using StarBEAST2 were still faster than gene tree relaxed clocks using *BEAST (Figure 4).

The increased performance of StarBEAST2 will enable researchers to analyse sequence data more quickly and disseminate their findings sooner; a large MCMC analysis which would currently take three months may now be performed in one week. In the case of phylogenomic data which has been subsetted for use with *BEAST, StarBEAST2 can be used to analyse more data for more precise estimates of species trees and other parameters in the same amount of time as a more limited *BEAST analysis.

### 3.6 Species tree branch length coverage and accuracy

Bayesian methods like StarBEAST2 produce both point estimates and credible intervals of inferred parameters. Ideally the point estimates will have low error, and the credible intervals will cover the corresponding true values. For well calibrated Bayesian methods, a 95% credible interval will include the true value 95% of the time.

Using a species tree relaxed clock with StarBEAST2 improved the coverage of branch length credible intervals (Figure 5A,B), but even when using a strict clock most simulated tip and internal branch lengths were within the corresponding credible intervals. This suggests that a strict clock model may be sufficient for studies using StarBEAST2 where substitution rate variation is not of direct interest. When using a strict clock with concatenation most internal branch lengths were outside the credible interval, but when using a relaxed clock were usually within the credible interval (Figure 5B). However even when using a relaxed clock, tip branch lengths were usually outside the credible intervals inferred by concatenation (Figure 5A).

**Figure 5:**
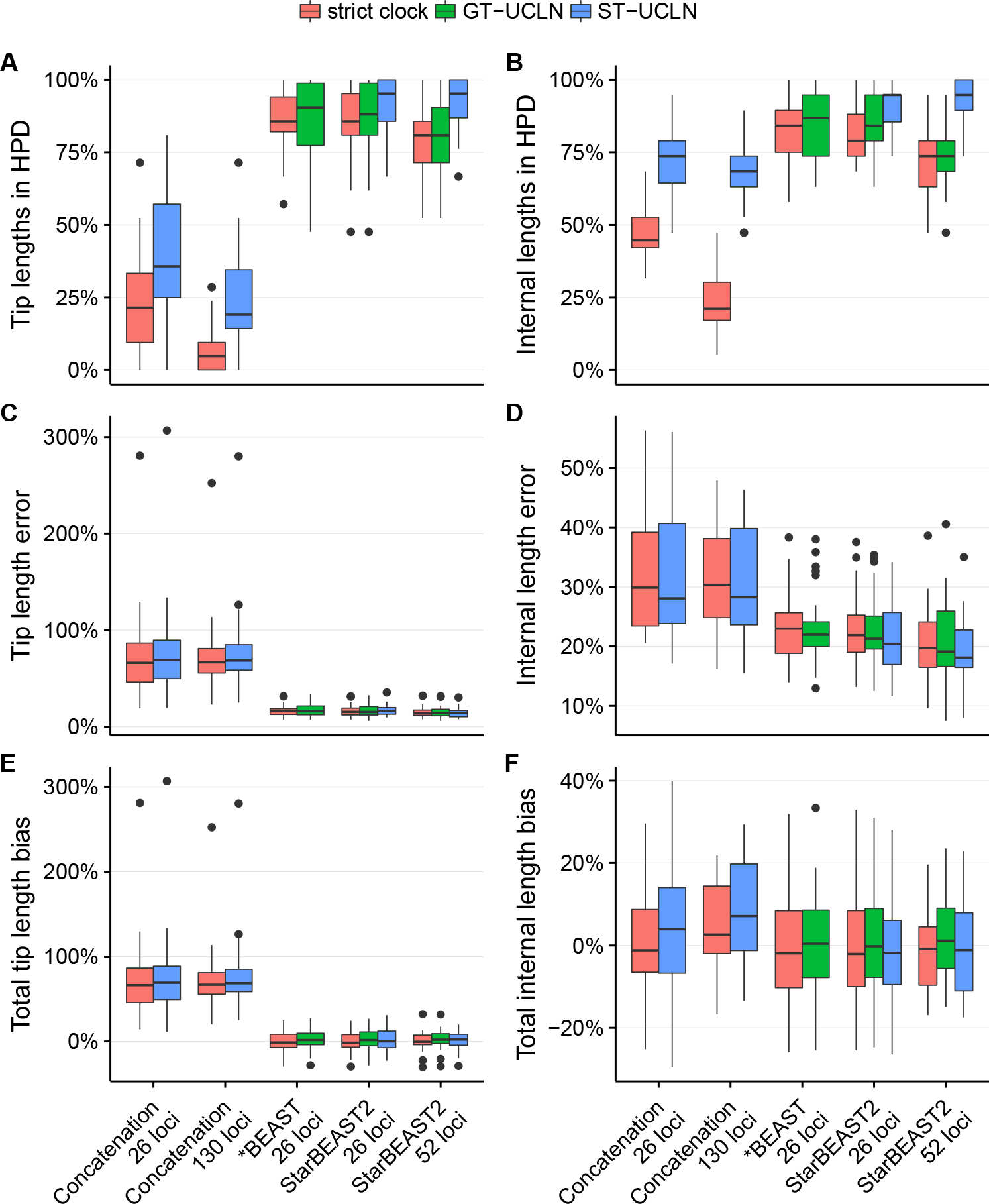
Coverage and accuracy of species branch lengths using different methods. Methods are StarBEAST2, *BEAST and BEAST concatenation with uncorrelated log-normal relaxed clocks applied to each gene tree (GT-UCLN) or to the species tree (ST-UCLN). (A,B) The percentages of true branch lengths present within the corresponding 95% highest posterior density (HPD) credible intervals. (C,D) The difference between the sum of estimated branch lengths and the sum of true branch lengths as a percentage of the sum of true branch lengths. (E,F) The sum of absolute differences between estimated and simulated branch lengths as a percentage of true tree length. N = 30.

Point estimates made by *BEAST and StarBEAST2 of both tip and internal branch lengths were more accurate than those made by concatenation (Figure 5C,D). The inaccuracy of tip branch lengths inferred using concatenation was driven by a strong bias towards overestimating tip branch lengths. For some replicates the sum of estimated tip branch lengths was more than double the sum of simulated tip branches lengths (Figure 5E). Relatively little overestimation of internal branch lengths was observed when using concatenation (Figure 5F).

Biased tip branch lengths are important because many published phylogenies show evidence of a slowdown in diversification rate (Moen and Morlon, 2014). If the ages of extant species are overestimated, this will artificially reduce the number of recent speciation events, mimicking a slowdown. We suggest that accurate inference of changing diversification rates requires species trees inferred by fully Bayesian MSC methods like StarBEAST2.

Using unphased sequences with ambiguity codes for heterozygous sites improved the accuracy of concatenation by reducing the bias in tip lengths to less than 40% (Figure S10). Ambiguity codes are treated by most phylogenetic methods (including BEAST) as base call uncertainty, indicating the nucleotide at a given site could be one of several possibilities. When used with unphased sequences, they actually indicate the presence of two nucleotides simultaneously, which is therefore a model violation. Using concatenation to analyse unlinked loci is also a model violation, but in the region of parameter space investigated by this simulation study the two errors may *partially* cancel out.

### 3.7 Species tree topology coverage and accuracy

We measured the coverage of species tree topologies and used the rooted Robinson-Foulds distance metric to measure the error associated with the maximum clade credibility (MCC) topology point estimates. As with branch lengths, using a relaxed clock with concatenation or a species tree relaxed clock with StarBEAST2 improved coverage. Regardless of clock model the coverage of concatenation was low; in less than 50% of replicates was the simulated topology in the credible set (Figure 6A).

**Figure 6:**
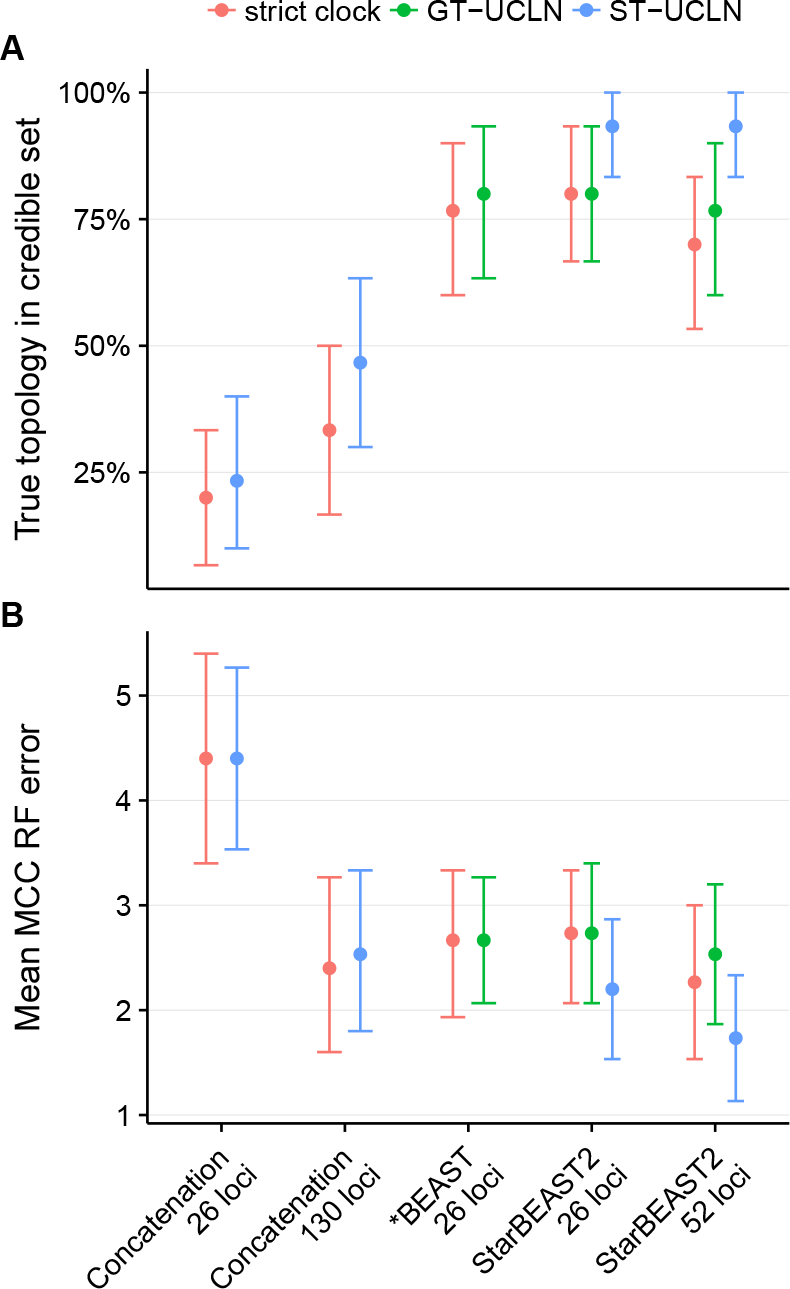
Coverage and accuracy of species tree topologies using different methods. Methods are StarBEAST2, *BEAST and BEAST concatenation with uncorrelated log-normal relaxed clocks applied to each gene tree (GT-UCLN) or to the species tree (ST-UCLN). (A) The percentage of true species tree topologies within the 95% credible set of topologies. (B) The average rooted Robinson-Foulds (RF) distance between the maximum clade credibility (MCC) species tree topology and the simulated true topology. Error bars are 95% confidence intervals calculated by bootstrapping. N = 30.

In terms of error rates, using 130 loci was similar to StarBEAST2 using 26 loci (Figure 6B). Using species tree relaxed clocks with StarBEAST2 was slightly more accurate than using strict clocks, but relaxed clocks did not improve the accuracy of concatenation (Figure 6B). Unlike branch lengths, topological accuracy was not improved by using un-phased sequences (Figure S11).

### 3.8 StarBEAST2 is superior at inferring substitution rates

While the convergence of species tree relaxed clock analyses took longer than for gene tree relaxed clocks in StarBEAST2, species tree relaxed clocks enable inference of species branch rates within an MSC framework. To gauge the accuracy of estimated branch rates, we used simple linear regressions with the true rate of each simulated branch as the explanatory variable, and the posterior expectation of the rate of that branch (conditional on the corresponding clade being monophyletic in the posterior samples) as the response variable. If all estimates are equally proportional to the truth, then the *R*^2^ coefficient of determination will equal 1. There are intrinsic limits to our ability to estimate substitution rates, namely that branch length is confounded with substitution rate (Thorne and Kishino, 2002).

For analyses of simulated data using 26 loci the *R*^2^ using StarBEAST2 was 0.39 and by doubling the number of loci to 52 was increased to 0.43. In contrast the *R*^2^ when using concatenation with 26 loci was 0.26 and even after increasing the number of loci to 130 it was only 0.33, in either case worse than StarBEAST2 using 26 loci (Figure 7). StarBEAST2 is clearly superior to concatenation at inferring branch rates.

**Figure 7:**
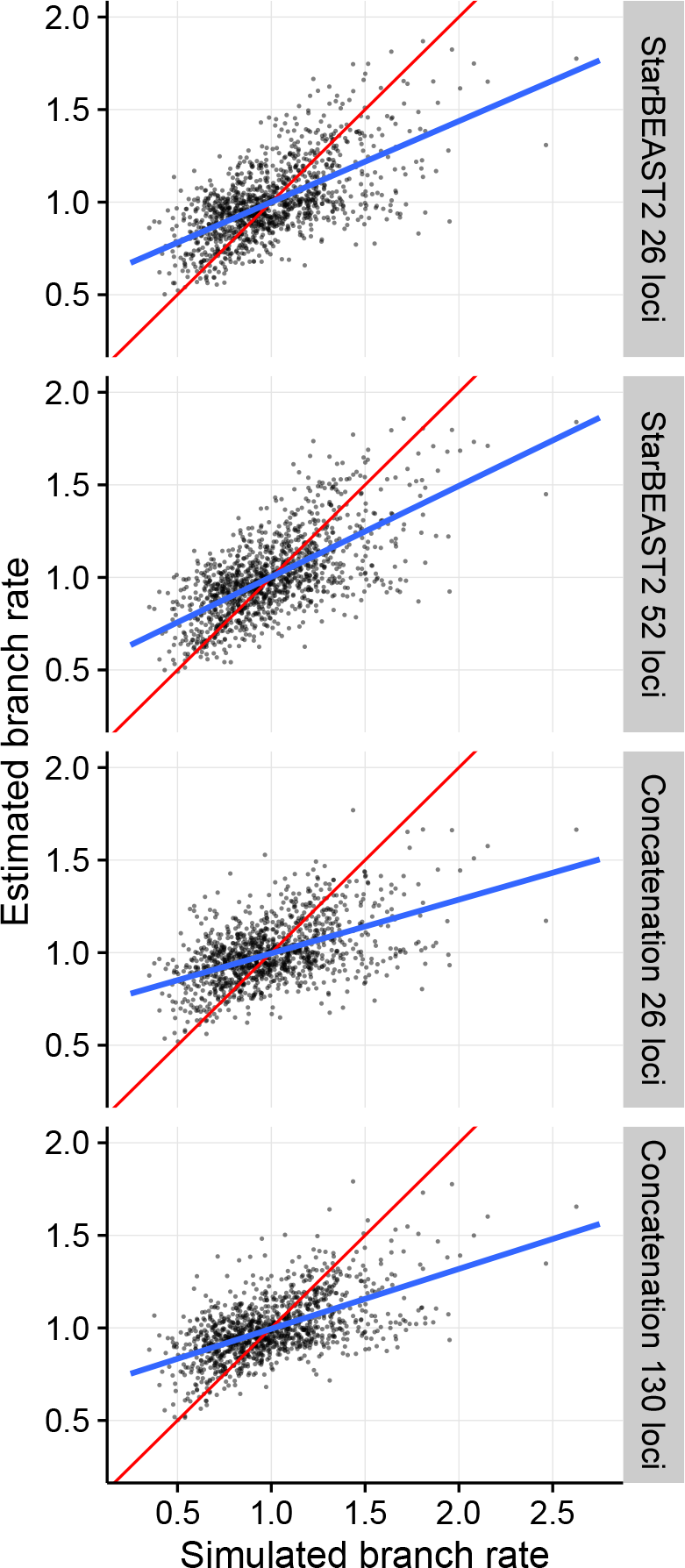
Estimates of species tree branch rates using BEAST concatenation versus Star-BEAST2. Estimated rates are the posterior expectations of each branch rate from each replicate. Root branch rates, which were fixed at 1, were excluded. In blue are simple linear regression lines of best fit, and in red are the *y = x* lines showing a perfect relationship between estimates and truth. N = 30.

Concatenation is an even worse estimator of branch rates when using unphased sequences with ambiguity codes for heterozygous sites. When applying concatenation to either 26 or 130 loci, *R*^2^ was very weak at 0.12 regardless of the number of loci (Figure S12).

## 4 Conclusions

When estimating divergence dates and substitution rates, the choice is often between using a subset of available loci with a fully Bayesian MSC method, or all available loci with concatenation. Researchers have often opted for the second choice, but we have shown that concatenation may not accurately estimate species ages or per-species substitution rates, even for trees of intermediate branch lengths. The increased performance of StarBEAST2 should further encourage the adoption of fully Bayesian MSC methods for estimating divergence times, and the new species tree relaxed clock will enable accurate inference of species branch rates despite ILS. StarBEAST2 is free and open source software; source code, development history and multiple tutorials are available on GitHub (https://github.com/genomescale/starbeast2).

## 5 Materials and Methods

For all StarBEAST2, *BEAST and concatenation analyses, the version of BEAST used was 2.4.4. For all simulations, the version of biopy (Heled, 2013) used was 0.1.9. Scripts used to perform all analyses will be available on GitHub (https://github.com/genomescale/starbeast2-manuscript/tree/master/scripts).

### 5.1 Mathematical correctness of StarBEAST2

Simulated trees were generated using biopy, and trees sampled from a prior distribution were generated using StarBEAST2 with all new features enabled. This included analytical integration of population sizes, coordinated tree topology and node height changing operators, and a species tree relaxed clock. 100,000 species trees were simulated, one gene tree was simulated per species tree with a rate of 0.5, and a second gene tree was simulated per species tree with a rate of 2.0.

100,000 species trees, 100,000 half rate gene trees and 100,000 double rate gene trees were sampled from the prior at a rate of one every 1000 after a 10% burn-in period.

Identical parameters were used for the simulation and for the StarBEAST2 run including the prior distributions. We fixed the number of species at 5 and the number of sampled haplotypes per species at 1. The birth and death rates were fixed at 200 and 100 per substitution respectively. Haploid population sizes followed an inverse gamma distribution with shape *α* = 3 and scale *β* = 0.004.

This procedure was repeated for both UCLN and for UCED species branch rates. Branch rates were sampled from a lognormal or exponential distribution, in either case with a mean of 1, discretised into 100 bins.

### 5.2 Reanalysis of *Pseudacris* sequence data

Phased and aligned *Pseudacris* sequence data were retrieved from Dryad (http://dx.doi.org/10.5061/dryad.23rc0). Replicating the original analysis we applied the HKY nucleotide substitution model (Hasegawa *et al*., 1985) to 22 out of 26 nuclear loci and the GTR model (Tavaré, 1986) to the remaining 4. For all models we used four discrete gamma categories to accommodate among-site rate variation (Yang, 1994). Transition/transversion rates and ratios and rate variation shape parameters were estimated, and empirical base frequencies used, all separately for each locus. The relative substitution rate of each locus was estimated using a lognormal prior with a mean *μ* in real space of 1 and a standard deviation σ of 0.6. We used a single haplotype sequence per individual per locus, halving the total number of sampled sequences to avoid wasting computational resources.

For inference of *Pseudacris* trees, we ran 30 independent StarBEAST2 chains for all 24 conditions for a total of 720 chains. The conditions were each possible combination of strict, species tree relaxed or gene tree relaxed clocks, analytical or MCMC population size integration, coordinated or naïve topology changing operators, and the inclusion or exclusion of coordinated height changing operators. Each chain used the same sequence data but was a partially independent estimate of convergence because a different random seed was used to initialise each chain.

A birth-death prior was used for the species tree and both the net diversification and extinction fraction hyperparameters were estimated. A gamma prior was used for MCMC estimated population sizes with a shape fixed at 2 and an estimated mean population size hyperparameter, matching the original *BEAST model (Heled and Drummond, 2010). The number of branch rate categories was equal to the number of estimated branch rates (as is the default in BEAST 2), and the standard deviation of the UCLN clock model was fixed at 0.3.

To ensure convergence of all chains, we ran each chain for an initial length of 2^24^ = 16,777,216 states, sampling every 2^11^ = 2,048 states. Initial chain lengths and sampling rates for all other analyses are in Table S8. ESS values were computed for all recorded statistics after discarding 12.5% of state samples as burn-in. Recorded statistics included (1) the posterior probability, (2) the coalescent probabilities of gene trees, (3) the overall prior probability, (4) the birth death prior probability of the species tree, (5) the phylogenetic likelihood, (6) the net diversification rate, (7) the extinction fraction, (8) the mean population size, and (9) the height of the species tree.

If any recorded statistic had an ESS below 200, the chain was resumed until the length of the chain had doubled. ESS values were then re-evaluated, again after discarding 12.5% of state samples. The length of a chain was continually doubled and ESS values re-evaluated until the ESS values of all recorded statistics were above 200. The rate at which trees and statistics were sampled was halved with every chain doubling so that the total number of samples remained constant. Two *BEAST GT-UCLN chains still had insufficient ESS values after running for 2^34^ = 17,179,869,184 states, but all other chains had converged. Estimated ESS values for all chains were used for analyses of computational performance.

ESS per hour was calculated by dividing the final ESS value for a given statistic by 87.5% of the total CPU time used by that chain to account for burn-in. Likewise ESS per million states was calculated by dividing the final ESS value by 87.5% of the total number of the states in the chain, then multiplied by one million. For all analyses of computational performance including graphs and linear models, the ESS rate for any given chain was that of the slowest converging statistic for that particular chain.

Average branch length in coalescent units was calculated by concatenating the output (after discarding the first 12.5% of states as burn-in from each chain) of all 30 chains which used the combination of MCMC population size integration, naïve topology operators, coordinated node height operators and species tree branch rates. For every sample in the combined posterior distribution, the coalescent length of each branch *τ*(2*N_e_*)^−1^ was calculated from its length in substitution units τ and its effective population size *N*_*e*_. The mean coalescent length of all branches across all samples was taken as the average.

### 5.3 Testing the effects of SPILS on estimated substitution rates

To test how SPILS affected estimates of per-species branch substitution rates, 96 fully asymmetric species trees were simulated with the topology ((((A,B),C),D),E). All species trees were simulated according to a pure birth Yule process (Yule, 1925) with a speciation rate of 10 per substitution.

Haploid population sizes for each branch were chosen independently from an inverse gamma distribution with a shape of 3 and a scale of 0.2. 100 gene trees with one individual per extant species were then simulated for each species tree according to the MSC process using biopy. Finally 1000nt sequence alignments were then simulated for each gene tree according to the Jukes-Cantor substitution model (Jukes and Cantor, 1969), equal base frequencies, no among-site rate variation, a strict molecular clock, and a substitution rate of 1 for each locus. Sequence alignments were simulated using Seq-Gen (Rambaut and Grassly, 1997).

BEAST concatenation and StarBEAST2 were then used to estimate the branch rates and divergence times with the species tree topology fixed to the truth. The same substitution model used for simulating sequences (i.e. Jukes-Cantor, no rate variation among sites or loci) was also used for inference. UCLN relaxed clocks were applied to the tree inferred by concatenation and to the StarBEAST2 species tree.

The same strategy as applied to *Pseudacris* was used to ensure convergence of Star-BEAST2, but for concatenation mean population sizes and coalescent probabilities are not part of the model and so were not recorded.

For every converged chain, the posterior expectation and 95% credibility intervals of per-species branch rates were calculated using the TreeAnnotator program supplied with BEAST.

### 5.4 Simulations to measure computational and statistical performance

All simulation parameters were chosen to be broadly similar to those observed in or estimated from the *Pseudacris* data set.

First, 30 species trees were simulated according to a birth-death process (Gernhard, 2008) using biopy with 21 extant species, a speciation rate of 100 and a death rate of 30. This corresponds to a net diversification rate of 70 and an extinction fraction of 0.3. Haploid population sizes for each branch were chosen independently from a gamma distribution with a shape of 2 and a scale of 0.002. For a species with annual generation times, as is the case for at least some *Pseudacris* species (Caldwell, 1987), and a substitution rate of 10^−9^ per year this corresponds to an effective population size Ne of around 2 million individuals per generation. Species branch rates were chosen from a log-normal distribution with a mean in real space of 1 and a standard deviation of 0.3, then scaled so that the mean of the branch rates for a given species tree was exactly 1. This ensured that per-branch rates always reflected relative differences in substitution rates.

For each species tree, 130 gene trees with two sampled haplotype sequences per species were simulated according to the MSC process using biopy. The mean clock rate for each locus was chosen from a log-normal distribution with a mean in real space of 1 and a standard deviation of 0.6.

For each gene tree, 600nt long sequence alignments were simulated using Seq-Gen (Rambaut and Grassly, 1997). An HKY model was used for all sequence alignments with equal base frequencies, a κ value of 3, and a four rate category discretised gamma model of among-site rate variation with a shape α value of 0.2. Hence all inference based on simulated data applied the HKY+Γ substitution model to all loci.

The same combinations of clock models, population size integration and new operators were explored using the simulated data as for *Pseudacris* to provide more generally applicable results regarding those new techniques. The same number of loci, convergence strategy and calculations of ESS rates were used for both. Both haplotype sequences were used for each species for *BEAST and StarBEAST2. One concatenation chain using phased haplotypes and a relaxed clock still had insufficient ESS values after running for 2^33^ = 8,589,934,592 states, but all other chains had converged. Estimated ESS values for all chains were used for analyses of computational performance and statistical coverage and accuracy.

### 5.5 Comparison of StarBEAST2 with *BEAST and concatenation

To compare the performance of StarBEAST2 with *BEAST, we ran 30 strict clock and 30 gene tree relaxed clock replicates of the *Pseudacris* reanalysis using the *BEAST package built into BEAST 2. We also reran each simulation replicate using *BEAST with a strict clock and gene tree relaxed clocks. The same priors, substitution models, and convergence strategies as used for StarBEAST2 were used with *BEAST.

For both data sets we reused the StarBEAST2 results for the combination of analytical population size integration, coordinated height-changing operators and naïve topology operators, which are all enabled by default in StarBEAST2. To demonstrate the scaling of StarBEAST2, we also reran each simulation replicate with an additional 26 loci (for a total of 52 loci) for all three clock models.

To compare concatenation with StarBEAST2, we reran each simulation replicate for each combination of either unphased ambiguity coded sequences or a single haplotype sequence per species, either a strict clock or species tree relaxed clock, and either the original 26 loci or with an additional 104 loci (for a total of 130 loci). We estimated the per-locus rates in the same way as for StarBEAST2, and applied the same convergence strategy as for SPILS concatenation. For species tree clock rates we used the same UCLN parameters as StarBEAST2 but applied to the concatenated tree, a model equivalent to that described by Rasmussen and Kellis (2007).

We also generated 30 replicates from a UCE data set of *Crocidura* shrews to show that StarBEAST2 can scale to 100 loci. For 1020 out of 1112 loci, the best fitting substitution model was either HKY or a nested model (Giarla and Esselstyn, 2015). To simplify configuring substitution models, we chose 100 unique loci at random, and separately for each replicate, from the set of loci which best fit HKY or a nested model. For each replicate we ran *BEAST with a strict clock or gene tree relaxed clock and a subset of 50 loci, StarBEAST2 with all three clock models and the same subset, and StarBEAST2 with all three clock models and all 100 loci. The same priors, substitution model and convergence strategies were used as for the simulated data set. All *Crocidura* MCMC chains converged.

### 5.6 Measurements of species tree coverage and accuracy

Branch length error is defined 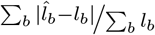 where *l_b_* is the true simulated branch length, and 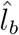 is the point estimate of the branch length, for a given species tree branch *b* in a set of branch lengths *B*. Total branch length bias is defined as 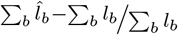. In this study, *B* is either the set of tip branches, or the set of internal branches excluding the root branch. Point estimates of branch lengths were calculated using the common ancestor method conditioned on the true simulated topology (Heled and Bouckaert, 2013). Highest posterior density regions were used for all credible intervals.

In this study, the rooted Robinson-Foulds distance (Robinson and Foulds, 1981) is the number of clades present in only one of the true tree *T*_1_ or the maximum clade credibility tree *T*_2_. Each 95% credible set of tree topologies was selected from topologies present in a given posterior sample, in order from high to low posterior probability, until the cumulative probability reached or exceeded 95%.

## 6 Supplementary Material

Supplementary figures and tables will be made available online.

## 7 Acknowledgments

This work was supported by a Rutherford Discovery Fellowship awarded to A.J.D. by the Royal Society of New Zealand. H.A.O. was supported by an Australian Laureate Fellowship awarded to Craig Moritz by the Australian Research Council (FL110100104). This research was undertaken with the assistance of resources from the National Computational Infrastructure (NCI), which is supported by the Australian Government. We wish to thank Jason Bragg and Renee Catullo for testing StarBEAST2 before its official release, Timothy Vaughan for suggesting the addition of a root height changing operator, Joseph Heled for insight into the multispecies coalescent, and Graham Jones for input regarding operator performance. We also thank Tanja Stadler for hosting H.A.O. and A.J.D. during part of the development of StarBEAST2, and thank Fábio Mendes, Matthew Hahn, Diego Mallo, two anonymous reviewers and the editor for suggesting valuable improvements to this manuscript.

